# Contrastive learning for antibody-antigen sequence-to-specificity prediction

**DOI:** 10.64898/2026.02.25.707916

**Authors:** Hyunho Lee, Karla Castro, Sean Renwick, Lucas Stalder, Wiona Glänzer, Rachita Kumar, NingNing Chen, Andreas Scheck, Alex Yermanos, Derek Mason, Sai T. Reddy

**Author notes:** Equal contribution (Hyunho Lee, Karla Castro, Sean Renwick).

## Abstract

Predicting which antibodies bind to which antigens directly from primary amino acid sequences remains a major challenge, as no current method can reliably determine this specificity at both a repertoire and proteome scale. Structure-based protein design frameworks can propose antibody binders to specified antigenic epitopes, but they do not solve the “sequence-to-specificity” task of mapping antibodies to cognate epitopes, and vice versa. Here, we introduce CALM (Cross-attention Adaptive Immune Receptor–Antigen Language Model), a dual-encoder plus cross-attentive decoder architecture that treats antibody–antigen recognition as molecular translation. Using contrastive learning, antigen and antibody encoders learn a shared embedding space that aligns cognate epitope–paratope binding pairs. CALM-1.0 is trained and evaluated on 4,138 curated antibody-antigen pairs obtained from the PDB-derived structural antibody database (SAbDab). On a leakage-controlled test split drawn from sequence clusters at 80% identity and unseen during training, CALM-1.0 achieves a mean top-1 retrieval (R@1) of 7%, with consistent performance across both directions (Ab→Ag and Ag→Ab). CALM establishes a foundation for bidirectional antibody-antigen sequence-to-specificity prediction with the potential to unify retrieval and generative design.

## INTRODUCTION

Deciphering the rules that govern antibody–antigen (Ab–Ag) specificity directly from primary amino acid sequence remains an enduring challenge at the intersection of biotechnology and immunology. The ambitious goal is to computationally predict, from sequence alone, which antibodies will bind to which antigens with high precision, a capability that would transform both therapeutic discovery and diagnostics. This “sequence-to-specificity” problem is the critical bottleneck in modern biologics drug discovery and immune repertoire profiling.

Foundational technologies such as hybridoma^1^ initiated the industrialization of antibody specificity, followed by combinatorial in vitro approaches like phage and yeast display,^2,3^ which together catalyzed the development of monoclonal antibodies into a cornerstone of modern therapeutics. Yet despite decades of progress, these paradigms remain inherently laborious and resource-intensive. A robust computational solution, such as an Immune Specificity Foundation Model (ISFM), would constitute a paradigm shift, unlocking two distinct but equally transformative capabilities. It would enable the forward design of novel de novo therapeutics on demand, from monoclonal antibodies to engineered T-cell receptors. Concurrently, an ISFM would solve the reverse interpretation problem, finally allowing us to “read” the vast, high-throughput immune repertoires of patients and deploy them as high-resolution diagnostics for oncology, autoimmunity, and infectious disease.

Despite the extraordinary progress of modern machine learning in protein biology, from accurate folding prediction^4^ to structure-based design^5^, an ISFM is elusive. The specificity of Ab–Ag recognition remains beyond what current models can reliably achieve. On the sequence side, protein language models (PLMs) have begun to approximate this grammar of structure and function. Foundational sequence-only PLMs such as ESM-2^6^ and ProGen^7^ learn representations predictive of structural and biophysical features.^8^ Within the antibody domain, specialized PLMs (e.g., AntiBERTy, IgLM, IgBERT/IgT5) have been developed to capture the intricacies of immune repertoire diversity, affinity maturation, and realistic antibody generation.^9–15^ At the structure end, diffusion-based co-folding frameworks such as AlphaFold 3^16^ and Boltz-2^17^ can now generate plausible Ab–Ag complexes and report pose-confidence metrics that correlate with binding in many regimes. Complementary backpropagation-through-structure methods extend this capability toward design: RFdiffusion/RFAntibody,^18^ Chai-2, Germinal, mBER, GeoFlow and BoltzGen families combine geometric diffusion, flow-matching, or gradient guidance to iteratively optimize sequence and structure together.^19–22^

Together, these advances underscore both the promise and the gap. While structure-based generative systems and large protein language models have transformed our ability to model proteins, no existing framework unifies antibody and antigen sequences into a single, scalable, sequence-native system that directly learns binding specificity in both directions. While contrastive co-embedding of antibody and antigen representations has recently been explored, for example, in EAGLE^23^, which incorporates a CLIP-like^24^ objective to align antibody sequences and antigen structures prior to diffusion-based generation, such approaches are embedded within structure-conditioned generative pipelines rather than developed as standalone, scalable retrieval models of binding specificity. A true Immune Specificity Foundation Model must operate at repertoire scale, generalize beyond seen antigens, and support both retrieval and conditional design without requiring structural inference at deployment. In this study we introduce CALM (Cross-attention Adaptive Immune Receptor–Antigen Language Model), a dual-encoder contrastive architecture that aligns antibody and antigen sequences in a shared embedding space. By training directly on paired binding data with leakage-controlled splits, CALM establishes a sequence-only foundation for bidirectional specificity prediction and provides a principled stepping stone toward a full ISFM capable of unifying discriminative alignment and generative translation within a single framework.

## METHODS

### Architecture of Cross-attention Adaptive Immune Receptor–Antigen Language Model (CALM)

Ab–Ag recognition fundamentally depends on sequence-level patterns that encapsulate both the structural grammar of antibodies—such as germline frameworks and CDR loop motifs—and the complex chemistry of epitopes. CALM is designed as a sequence-native model that learns a shared latent space for antibodies and antigens and can, in principle, support conditional generation of partner sequences (**Fig. 1**). The core of the present work is a dual-encoder co-embedding module: modality-specific encoders for antibody (paratope) and antigen (epitope) sequences are trained with a contrastive objective to place true binder pairs close together in embedding space while pushing non-binders apart. This stage is fully implemented and forms the basis of all experiments reported here.

**Fig 1.**
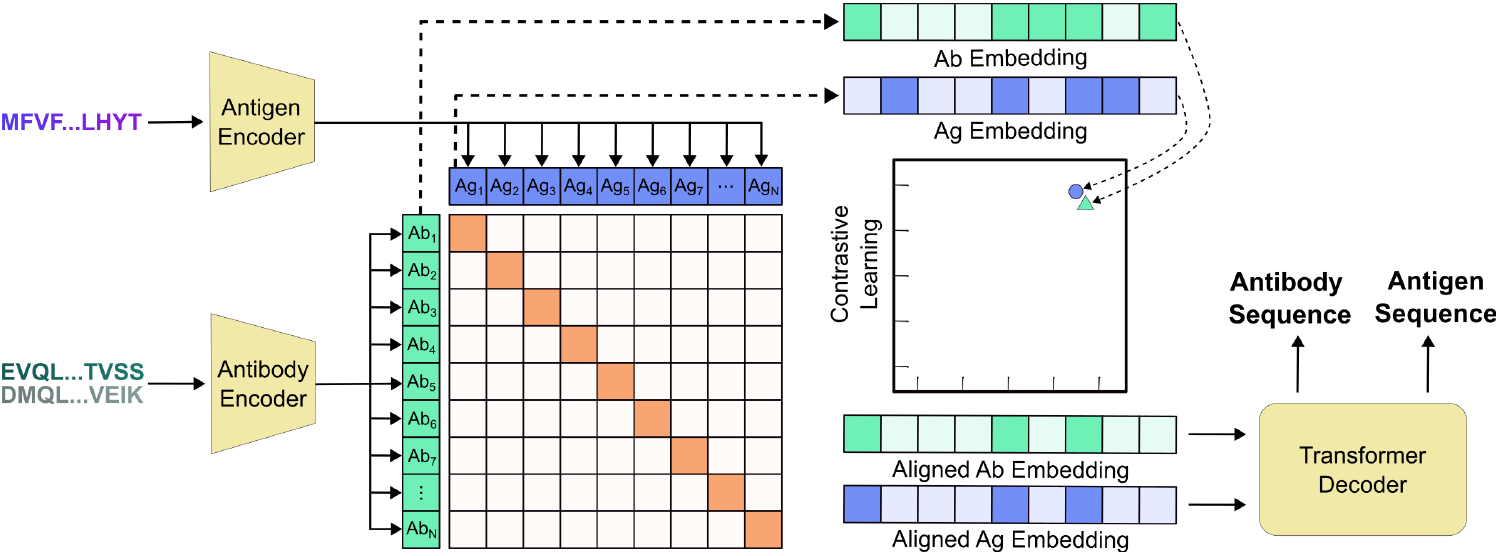
Schematic of CALM (Cross-attention Adaptive Immune Receptor–Antigen Language Model) architecture. CALM is a sequence-based architecture that addresses both the Ab–Ag binding specificity prediction task and the binder generation task using contrastive learning and a generative model. CALM projects Ab and Ag sequences into a joint embedding space and maximizes the contrastive alignment of the embeddings between binding Ab–Ag pairs. The jointly aligned embeddings are passed to an Encoder-Decoder Transformer for de novo sequence generation. The Transformer attends to the aligned Ag embedding to generate novel Ab sequences with binding specificity, and simultaneously, attends to the aligned Ab embedding to generate novel Ag sequences.

CALM’s co-embedding objective is inspired by dual-encoder paradigms that align heterogeneous modalities via contrastive learning, most famously in vision-language models such as CLIP, where paired images and captions are pulled together in embedding space and negative pairs are pushed apart^24^. In CALM, antibody and antigen embeddings, initialized using AntiBERTy^10^ and ESM-2^6^, respectively, are aligned so that true binder pairs become geometrically proximal in a shared embedding space. This alignment supports retrieval tasks in both directions (Ab→Ag, Ag→Ab) and is the only component trained and evaluated in this study.

In addition to this co-embedding stage, we propose, but do not train or benchmark here, an autoregressive decoder with cross-attention as a natural extension of the architecture. Conceptually, this decoder would mirror encoder-decoder Transformers in neural machine translation^25–27^: queries from the decoder attend to keys/values produced by the antibody or antigen encoder to condition generation of the partner sequence (antibody→epitope or epitope→antibody), potentially with task-prefix tokens (“generate-antibody”, “generate-epitope”) akin to text-to-text models such as T5.^28^ We include this decoder design to illustrate how the same CALM backbone could eventually unify discriminative alignment and generative translation within one model, but all results in this manuscript are based solely on the Stage-1 dual-encoder contrastive training. Future work will be required to implement, train, and systematically evaluate the proposed decoder for epitope mapping and conditional antibody or antigen design.

### Data curation and pre-processing

4,941 antibody-antigen pairs were extracted from SAbDab^29^ together with their structure files and converted to a standardized CALM data schema (**Table 1** and **Fig. 2**). Antigens were required to be of type *protein* and duplicate pairs were dropped from the dataset. Variable heavy (VH) and light (VL) sequences were extracted from the provided antibody sequences using AbNumber^30^ following Chothia numbering. Single-chain variable fragment (scFv) entries were split into VH and VL chains with AbNumber and were treated identically to monoclonal antibodies. We restricted entries to Ab–Ag pairs with only a single antigen chain and required that at least one antibody chain (heavy or light) was present. The final dataset includes 3,251 entries containing both VH and VL sequences, 874 entries with only VH sequences, and 13 entries with only VL sequences. To facilitate efficient clustering and embedding of the retrieved sequences, several restrictions regarding sequence length were applied. Antigens were required to be between 19 and 550 amino acids long and variable regions longer than 141 amino acids were removed. The final dataset consists of 4,138 Ab–Ag pairs, containing 3,332 and 3,467 unique antibody and antigen sequences, respectively.

**Table 1.** Standardized CALM data format. All data collected was parsed to follow this CALM-schema. Data labels in bold are required for every data point. Binding complexes may be missing one antibody chain (VH or VL), but not both.

| Data labels | Description | Example |
| --- | --- | --- |
| **Unique_AG_VH_VL_Hash** | Unique identifier for complex | f3c1dbd8da8c1... |
| **Antibody_VH_ID,**<br>**Antibody_VL_ID, Antigen_ID** | Chain ID in reference PDB-file | “H”, “L”, “A” (can vary significantly) |
| **Antibody_VH_AA,**<br>**Antibody_VL_AA, Antigen_AA** | Amino acid sequence | ...SGTNGTKRFDNP... |
| Antibody_VH_Paratope,<br>Antibody_VL_Paratope,<br>Antigen_Epitope | Mask for each chain indicating where each residue is within 5Å | ...0000100111000... |
| VH_CDR{1-3}_Chothia_Mask,<br>VH_FW_Chothia_Mask,<br>VL_CDR{1-3}_Chothia_Mask,<br>VL_FW_Chothia | Mask for each antibody chain region | CDR1: ...0000111110000...<br>FW: ...111100001111... |
| Antigen_AA_Uniprot,<br>Antigen_Uniprot_ID | UniProt ID for antigen sequence, and associated amino acid sequence | Q07337, ...MTLFLFISCN... |
| Antibody_ScFv_ID,<br>Antibody_ScFv_AA | Chain ID in reference PDB-file of ScFv |  |
| **Binder_Status** | Binding (“Y”) or Non-binding (“N”) | “Y” |
| Antibody_Type | Type of antibody (Monoclonal, ScFv, etc.) | Monoclonal |
| **Cluster_id** | MMseqs2 cluster ID, used for train/test/validate split | “C000001” |

**Fig 2.**
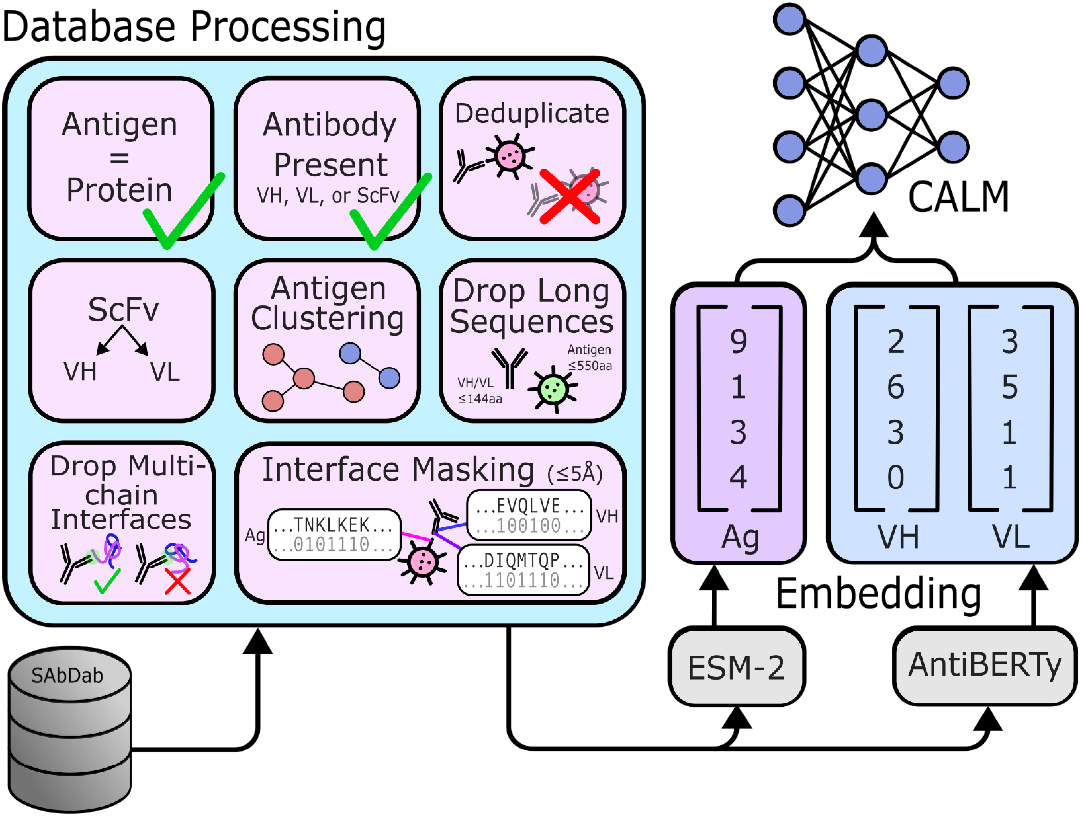
Dataset curation and preprocessing. Ab–Ag complex structures were extracted from SAbDab. Complexes were filtered to include only protein antigens and at least one antibody chain, with duplicate sequences removed. VH and VL regions were extracted from the provided sequences with AbNumber. ScFv sequences were split into VL and VH chains using AbNumber where possible. Sequences exceeding the embedding dimension limit (141 for VH and VL, and 550 for Antigens) and complexes with multi-chain antigen binding interfaces were excluded. Standardized Chothia masks were generated via AbNumber, along with epitope/paratope masks defined by residues within a 5 Å proximity at the binding interface, generated via BioPython. Complexes were clustered based on antigen sequence with MMseqs2 at 40%, 60%, and 80% sequence similarity before they were embedded with ESM-2 and AntiBERTy.

### Sequence embeddings

For all VH, VL, and antigen sequences, embeddings were generated by padding the sequences to their maximum length and tokenizing them with the tokenizers of the respective embedding models. Antibody sequences were embedded using the pre-trained antibody language model AntiBERTy.^10^ Both variable regions were embedded separately and then concatenated. Antigen sequences were embedded with the pre-trained ESM-2(model version esm2_t33_650M_UR50D^6^). For both antibody and antigen sequences, embeddings were taken from the last hidden layers. Token embeddings of start and end tokens were removed and embedding vectors at padded positions were replaced with zero-vectors.

### Epitope and paratope mask generation

Binary binding-site masks were generated for all data points with distance-based contact annotations from the associated structure files using BioPython. For each antibody chain, all residues within 5 Å of the antigen were indicated with “1” and all other residues were indicated with a “0” to build paratope masks. This process was repeated for each antigen, where all residues within 5 Å of either antibody chain were indicated with a “1” and all other residues were indicated with a “0” to build epitope masks.

### Clustering

To ensure leakage-controlled evaluation of the model during cross-validation, all Ab-Ag pairs were clustered based on their antigen amino acid sequence identity using MMseqs2^31^. Clustering was performed on antigen amino acid sequences with the command “mmseqs cluster … --min-seq-id {*training_sequence_identity*} -c 0.4 --cluster-mode 1 --max-seqs 10000000 --single-step-clustering -s 7.5”, with a *training_sequence_identity* of 40%, 60%, or 80%, depending on the dataset. An Unclustered (UC) dataset was also prepared without data stratification according to antigen sequence similarity. Instead, VH and VL sequences were clustered with the command “mmseqs cluster … --min-seq-id {*UC_sequence_identity*} -c 0.4 –max-seqs 10000000” with *UC_sequence_identity* of 90% or 95% (UC90, UC95) in order to evaluate in-distribution performance. Additionally, a Randomly Shuffled (RS) dataset was prepared where Ab-Ag pairs were randomly mismatched to provide a random baseline for model performance. To avoid potential bias from over-represented clusters in the final datasets used for training and validation (**Fig. S1**), we subsampled up to 25 Ab–Ag pairs per cluster, resulting in 2,809 and 2,981 Ab–Ag pairs for clustering at 40% and 60% antigen sequence identity, and 3,094 Ab–Ag pairs for clustering at 80%, UC90, UC95, and RS, respectively. We note that subsampling for the UC and RS datasets is based on antigen sequence clustering at 80%. Data splits based on this subsampled dataset were further validated to be leakage-safe by computing pairwise alignments for all entries using Parasail’s Needleman-Wunsch implementation^32^ in the case of antigen sequence clustering at 40%, 60%, and 80%. A similarity score for two sequences was computed as percent identity over the aligned region scaled by the alignment coverage. If any members from two different MMseqs2 clusters had a similarity score higher than 0.4, 0.6, or 0.8, respectively, the affected clusters were merged, such that the larger cluster absorbed the smaller cluster.

### CALM architecture and training

Stage-1 (contrastive co-embedding): CALM jointly trains the antigen encoder and antibody encoder to maximize the cosine similarity between the feature representations of Ab-Ag binding pairs. Starting from the pre-trained embeddings, epitope (Ep) and paratope (Pa) masks are applied to the antigen and antibody embeddings, respectively, allowing the model to focus on the residues directly involved in binding. Each encoder then uses a feed-forward network as a projection head to map the representations into a joint embedding space. The joint embeddings for antigen and antibody are then length-wise mean pooled to compute their cosine similarity. We use a symmetric multi-positive contrastive loss to handle cases where each epitope or paratope has multiple positive pairs within the batch (e.g. two different paratopes targeting the same epitope). Let *s*_*i,j*_ denote the similarity logits, *N* denotes batch size, *Y*_*i,j*_ ∈ {0, 1} denotes label, and the normalized probability distribution 𝒫_*i,j*_:

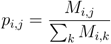

For each direction (Pa→Ep and Ep→Pa), we defined the multi-positive contrastive loss:

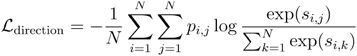

Then, the final loss is:

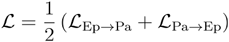

For training, we use the Adam optimizer with decoupled weight decay regularization and set the learning rate using cosine-annealing scheduler with warm restarts. We use a mini-batch size of 1,024 for training and 256 for validation and testing due to the low-data regime.

The model was trained using the AdamW optimizer with an initial learning rate of 0.001 and weight decay of 0.2. The learning rate was scheduled using a cosine annealing with warm restarts. The learnable temperature parameter τ was initialized to 0.2.

### Model evaluation

We performed nested 5-fold cross-validation with cluster-aware splitting, such that all Ab–Ag pairs from the same cluster were assigned to the same partition. In each outer fold, 20% of the clusters were held out as the test set, with the remaining clusters used for training and validation. Accordingly, this procedure yielded an approximate 65:15:20 train:validation:test ratio of Ab–Ag pairs per fold. Final numbers are reported as average across the five folds.

In standard CLIP-style retrieval, the ground-truth binding pair is treated as the sole positive example, while all the other entries in the mini-batch are considered negatives (in-batch negatives). However, due to the low-data regime, identical epitope or paratope sequences may appear multiple times across different Ab–Ag pairs within a single mini-batch. To address this, we adopted a multi-positive label in which all entries sharing the identical epitope or paratope sequences were treated as valid positives, even if they originated from different Ab–Ag pairs.

Model performance was evaluated using Recall@k (R@k) with *k* = 1, 5, 10 across different clustered datasets, i.e. at 40%, 60%, 80% antigen sequence identity, UC, and RS to represent a wide spectrum of out-of-distribution (OOD) and in-distribution performance. Under the multi-positive setting, Recall@k was computed using a binary success criterion per query, rather than normalizing by the total number of positives, as normalization by multiple redundant positive instances would artificially penalize the model. This formulation is equivalent to Hit@k, however, we refer to it as Recall@k for consistency.

## RESULTS

### CALM retrieval performance of antibody-antigen pairs

We evaluated CALM on 4,138 antibody–antigen (Ab–Ag) pairs derived from SAbDab using leakage-controlled antigen clustering at 40%, 60%, and 80% sequence identity thresholds. Because clustering threshold determines the degree of antigen uniqueness in the test set, lower identity cutoffs impose progressively stronger out-of-distribution (OOD) generalization challenges. Retrieval was assessed bidirectionally (Ag→Ab and Ab→Ag) using recall at top-k (R@1, R@5, R@10), with random shuffling serving as a baseline.

At the strictest OOD split (40% identity), CALM achieves R@1 of ∼2% and R@10 of ∼9% in both retrieval directions, exceeding the random baseline (R@1 ∼0.6%; R@10 ∼5.4%). Performance improves at 60% clustering, with R@1 increasing to ∼3% and R@10 to ∼12%. At 80% clustering, retrieval further increases to R@1 ∼6% and R@10 ∼16%, representing approximately a three-fold improvement over random at top-1 retrieval and more than double the random baseline at top-10. R@5 follows the same monotonic trend across clustering thresholds (**Fig. 3, Table 2**).

**Table 2.**
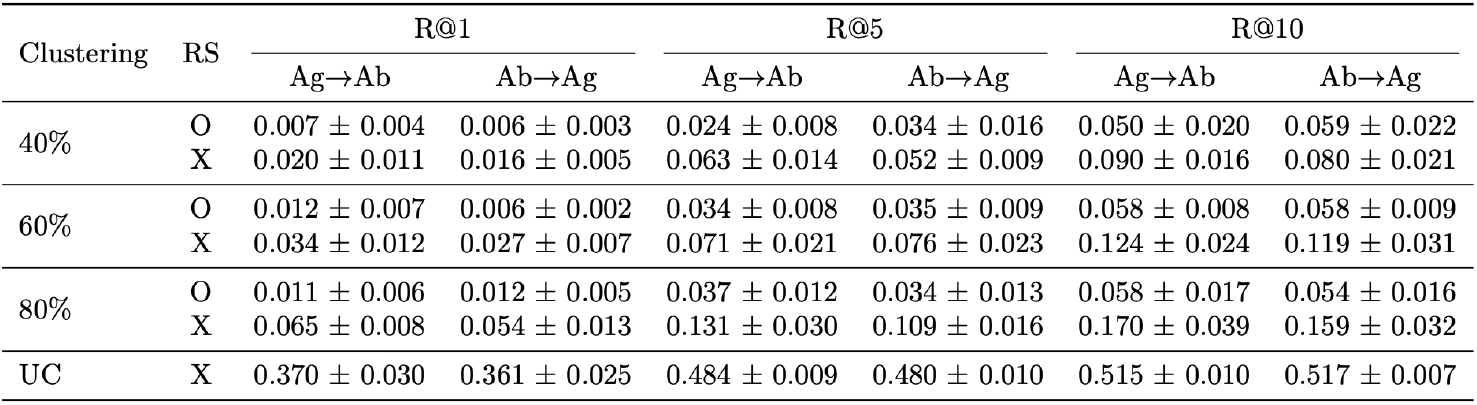
Retrieval performance of CALM on Ab–Ag pairs. Evaluation was conducted using full-length Ab–Ag sequences. Test sets (n=512 per fold) from five cross-validation folds were included. O and X denote with and without random shuffling (RS), respectively.

| Clustering | RS | R@1 |  | R@5 |  | R@10 |  |
| --- | --- | --- | --- | --- | --- | --- | --- |
|  |  | Ag→Ab | Ab→Ag | Ag→Ab | Ab→Ag | Ag→Ab | Ab→Ag |
| 40% | O | 0.007 ± 0.004 | 0.006 ± 0.003 | 0.024 ± 0.008 | 0.034 ± 0.016 | 0.050 ± 0.020 | 0.059 ± 0.022 |
|  | X | 0.020 ± 0.011 | 0.016 ± 0.005 | 0.063 ± 0.014 | 0.052 ± 0.009 | 0.090 ± 0.016 | 0.080 ± 0.021 |
| 60% | O | 0.012 ± 0.007 | 0.006 ± 0.002 | 0.034 ± 0.008 | 0.035 ± 0.009 | 0.058 ± 0.008 | 0.058 ± 0.009 |
|  | X | 0.034 ± 0.012 | 0.027 ± 0.007 | 0.071 ± 0.021 | 0.076 ± 0.023 | 0.124 ± 0.024 | 0.119 ± 0.031 |
| 80% | O | 0.011 ± 0.006 | 0.012 ± 0.005 | 0.037 ± 0.012 | 0.034 ± 0.013 | 0.058 ± 0.017 | 0.054 ± 0.016 |
|  | X | 0.065 ± 0.008 | 0.054 ± 0.013 | 0.131 ± 0.030 | 0.109 ± 0.016 | 0.170 ± 0.039 | 0.159 ± 0.032 |
| UC | X | 0.370 ± 0.030 | 0.361 ± 0.025 | 0.484 ± 0.009 | 0.480 ± 0.010 | 0.515 ± 0.010 | 0.517 ± 0.007 |

**Fig 3.**
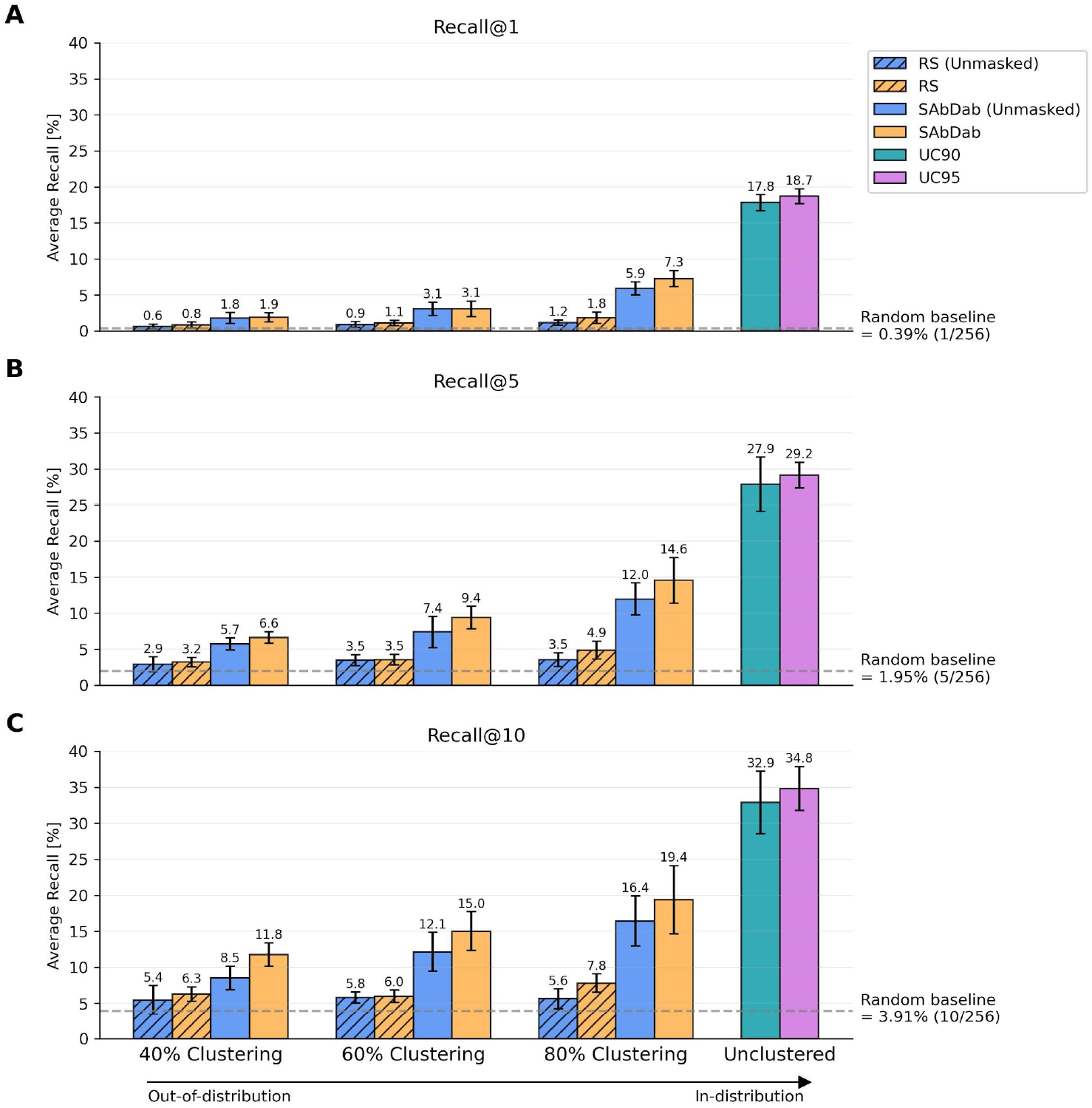
CALM In- and Out-of-distribution performance. CALM model performance reported as **A**. average Recall@1, **B**. average Recall@5, and **C**. average Recall@10, showing the mean of the Ab→Ag and Ag→Ab retrieval tasks. Performance is evaluated for a gradient of In- and Out-of-distribution test sets. Measured recall is contrasted to a randomly shuffled (RS) dataset (hatched) as well as the theoretical random retrieval success denoted as *Random baseline*. For antigen sequence clustering at 40%, 60%, and 80% and RS recall is shown with (orange) and without (blue) masking applied.

Across all clustering regimes, performance is consistent in both directions (Ag→Ab and Ab→Ag), with overlapping confidence intervals for R@k (**Table 2**). This directional symmetry indicates that CALM learns a balanced shared embedding space capable of retrieving cognate binding partners irrespective of query modality.

### CALM retrieval performance of paratope-epitope pairs

To assess whether restricting the model inputs to parameters corresponding to paratope-epitope binding interfaces enhances specificity learning, we retrained CALM using binary paratope and epitope masks derived from structural contact annotations. These masks limit the encoder inputs to residues within 5 Å of the binding interface, isolating local interaction determinants while excluding full-sequence context. Retrieval performance was evaluated under the same clustering regimes using R@k metrics.

Masked paratope–epitope retrieval exhibits a similar scaling pattern with distributional shift. At 40% clustering, CALM achieves R@1 of ∼2% and R@10 of ∼12%, exceeding the corresponding random baseline (R@1 ∼0.8%; R@10 ∼6.3%). At 60% clustering, performance increases to R@1 ∼3% and R@10 ∼15%. At 80% clustering, masked retrieval reaches R@1 ∼7% and R@10 ∼19%. As with full-sequence retrieval, R@5 increases consistently across clustering thresholds (**Fig. 3, Table 3**).

**Table 3.** Retrieval performance of CALM on Pa–Ep pairs. Evaluation was conducted using Pa–Ep sequences. Test sets (n=512 per fold) from five cross-validation folds were included. O and X denote with and without random shuffling (RS), respectively.

| Clustering | RS | R@1 |  | R@5 |  | R@10 |  |
| --- | --- | --- | --- | --- | --- | --- | --- |
|  |  | Ag→Ab | Ab→Ag | Ag→Ab | Ab→Ag | Ag→Ab | Ab→Ag |
| 40% | O | 0.007 ± 0.005 | 0.009 ± 0.005 | 0.030 ± 0.011 | 0.034 ± 0.008 | 0.059 ± 0.015 | 0.067 ± 0.013 |
|  | X | 0.020 ± 0.007 | 0.018 ± 0.007 | 0.066 ± 0.008 | 0.066 ± 0.009 | 0.119 ± 0.017 | 0.116 ± 0.017 |
| 60% | O | 0.011 ± 0.005 | 0.011 ± 0.004 | 0.032 ± 0.009 | 0.039 ± 0.007 | 0.054 ± 0.013 | 0.065 ± 0.009 |
|  | X | 0.033 ± 0.011 | 0.029 ± 0.012 | 0.088 ± 0.018 | 0.099 ± 0.017 | 0.148 ± 0.029 | 0.152 ± 0.026 |
| 80% | O | 0.018 ± 0.008 | 0.018 ± 0.008 | 0.047 ± 0.014 | 0.050 ± 0.011 | 0.078 ± 0.016 | 0.078 ± 0.012 |
|  | X | 0.074 ± 0.012 | 0.071 ± 0.012 | 0.145 ± 0.024 | 0.146 ± 0.040 | 0.193 ± 0.050 | 0.195 ± 0.046 |
| UC | X | 0.407 ± 0.021 | 0.404 ± 0.020 | 0.523 ± 0.023 | 0.518 ± 0.021 | 0.557 ± 0.023 | 0.552 ± 0.026 |

Direct comparison indicates that paratope–epitope masking yields systematically higher R@k values than full antibody–antigen retrieval at equivalent clustering thresholds. This suggests that restricting the model inputs to interface residues concentrates the binding signal and reduces sequence-level noise, improving ranking accuracy under OOD conditions.

### CALM retrieval performance of in-distribution paratope-epitope pairs

To evaluate CALM under an in-distribution regime with respect to antigens, we trained and evaluated a paratope–epitope (Pa–Ep) model without stratifying the data by antigen sequence similarity. Instead, to prevent trivial memorization of highly similar antibodies, we performed leakage-controlled clustering based on antibody sequence identity at 90% and 95% thresholds. Retrieval performance was assessed bidirectionally (Ep→Pa and Pa→Ep) using R@k metrics.

At 90% antibody clustering, CALM achieves R@1 of ∼18%, with R@5 of ∼28% and R@10 of 33% in both directions. Increasing the clustering threshold to 95% yields a modest additional improvement, with R@1 rising to ∼19% and R@10 to ∼35%. The close agreement between retrieval directions indicates that the shared embedding space maintains directional symmetry even under antibody-stratified splits (**Fig. 3, Table 4**).

**Table 4.** Retrieval performance of CALM on antibody-clustered data splits. Instead of clustering by antigen sequence identity, data splits based on VH and VL sequence identity (UC90 and UC95) were used to evaluate the model with antigen sequences remaining in-distribution while introducing controlled variation in antibody sequences. Test sets (n=512 per fold) from five cross-validation folds were included.

| Clustering | R@1 |  | R@5 |  | R@10 |  |
| --- | --- | --- | --- | --- | --- | --- |
|  | Ag→Ab | Ab→Ag | Ag→Ab | Ab→Ag | Ag→Ab | Ab→Ag |
| 90% | 0.178 ± 0.016 | 0.179 ± 0.011 | 0.277 ± 0.035 | 0.281 ± 0.041 | 0.329 ± 0.044 | 0.329 ± 0.043 |
| 95% | 0.188 ± 0.011 | 0.186 ± 0.013 | 0.291 ± 0.020 | 0.292 ± 0.017 | 0.348 ± 0.030 | 0.348 ± 0.032 |

Compared to the antigen-clustered OOD regime, these results show substantially higher retrieval accuracy, consistent with the model operating within a familiar antigen distribution while generalizing across antibody sequence variation. The relatively similar model performance observed between clustering at 90% and 95% further suggests that residual antibody sequence similarity modestly facilitates retrieval but does not dominate performance, indicating that CALM captures transferable interface-level determinants of binding beyond near-sequence identity.

## DISCUSSION

CALM is a sequence-native system trained with contrastive learning to align antibody and antigen sequences in a shared embedding space. It has the potential to be developed into an ISFM by supporting repertoire-scale analysis, antigen discovery, and antibody design. In this work we only report initial retrieval baselines on SAbDab-scale supervision. We focus on Stage-1 co-embedding and show that CALM achieves non-trivial top-k retrieval on leakage-controlled SAbDab splits where test antigens are held out by sequence identity. The decoder is specified architecturally but not trained or benchmarked here.

Current structure-conditioned systems address a different problem: antibody design given a defined epitope (Ep → Ab). RFdiffusion/RFAntibody, Chai-2, GeoFlow, Germinal, mBER, and BoltzGen report post-filter wet-lab hit rates in single- to double-digit percent range^17–22,33^, but these assay-level hit-rates after structural filtering are not directly comparable to CALM’s pool-dependent retrieval. CALM operates on paired VH–VL and antigen sequences, models both directions, and uniquely performs Ab→Ag or Pa→Ep prediction, a capability with no analogue in structure-first workflows.

To contextualize CALM’s performance, it is useful to relate our evaluation framework to established conventions in contrastive learning. For example, CLIP (Contrastive Language-Image Pre-training)^24^ is a vision-language model that learns to associate images with natural language descriptions by training two separate encoders, one for images and one for text, to place matching image-text pairs close together in a shared embedding space using the same contrastive objective (symmetric infoNCE over cosine similarities) that CALM employs. CLIP evaluates zero-shot transfer by presenting new images and classifying them against a fixed vocabulary of text prompts derived from known categories. CALM’s in distribution performance (no antigen clustering) is analogous: every test antibody sequence is unseen during training, and the task is to retrieve the correct binding partner from a pool of 256 candidates drawn from antigen clusters represented in the training set. This is a step towards zero-shot retrieval for antibodies where the model must generalize to antibody sequences not present in the training data. It is notable that CALM achieves R@1 of 17.8% and R@10 of 32.9% in this regime (approximately 46x and 8.4x above random, respectively) from ∼3,000 training pairs. In comparison, CLIP required 400 million image-text pairs to reach similar zero-shot transfer performance, and demonstrated that performance scales as a power law with data.^24^ While the tasks differ in domain complexity, both are cross-modal recognition problems learned via the same contrastive objective. The out-of-distribution (OOD) evaluations (80%, 60%, 40% antigen clustering) represent progressively harder tasks where test antigens share less than the specified sequence identity with training antigens. When trained with these stratified datasets, CALM maintains R@10 of 19.4% at 80% OOD and 11.8% at 40% OOD, indicating the embedding space captures generalizable features of binding specificity beyond memorization.

One framework for interpreting the anomalous data efficiency is the recent theory of computational convergence, which establishes that the softmax attention function of transformers is mathematically equivalent to the Boltzmann distribution governing antibody-antigen contact probabilities, and the InfoNCE contrastive objective is mathematically equivalent to the negative log of clonal selection probability^34^. If the mathematics governing immune recognition and the mathematics governing contrastive learning are identical, then a contrastive model on binding data does not require function approximation - discovering an unknown function from scratch. Instead it may resemble parameter estimation within a known equation, which is much more data efficient. Under this interpretation, each new paratope-epitope sequence samples a new region of the equation’s parameter space. The findings here raise the possibility that immunological scaling laws, governing how models scale when architecture and physics are mathematically aligned, may differ from the power-law neural scaling laws observed in established deep learning applications.^35^

### Limitations of Study

Several limitations should be noted. CALM reports only retrieval performance; the decoder is not benchmarked and no generation results are presented here. No wet-lab validation has been performed. Future directions include systematic scaling experiments to increase dataset diversity and size; decoder training for conditional antibody generation; incorporation of hard negatives into training; addition of immune repertoire training data; and wet-lab validation on in distribution and novel targets and epitopes.

## Acknowledgments

We would like to thank Damiano Sgarbossa for helpful discussions and for comments on the manuscript. We would like to acknowledge the Swiss National Supercomputing Centre (CSCS) for their support in computing time. In particular, we thank Lukas Drescher and Elia Palme for their technical support.

## Academic and non-profit research

We will release the CALM-1.0 research prototype (training/inference code, configs, evaluation notebooks), along with processed SAbDab splits and full reproduction scripts for the results reported here.

## Commercial use & IP notice

Use of the CALM methods, architecture, or trained weights in commercial settings requires a license. BIIE and ETH Zürich have filed patent applications covering methods and systems related to CALM and related rights are reserved. Commercial licensing inquiries may be directed to BIIE/ETH Zürich technology transfer.

## SUPPLEMENTARY FIGURES

**Fig S1.**
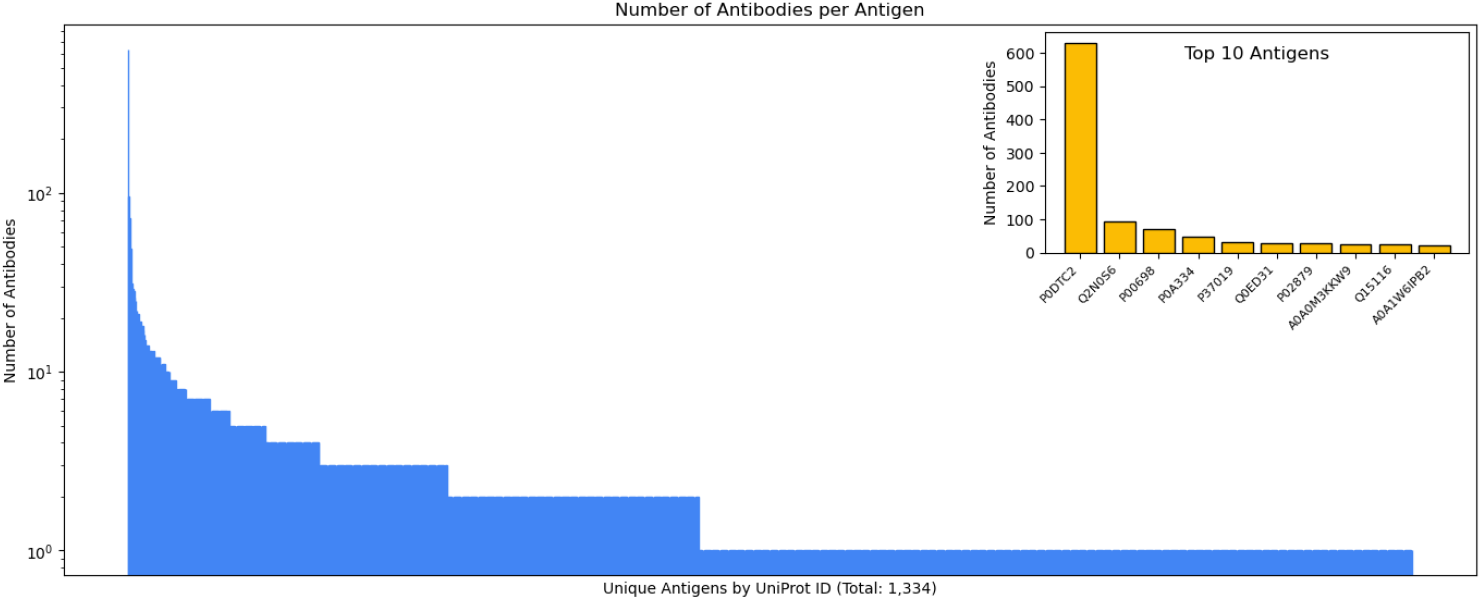
Number of antibodies per antigen. Distribution of the number of antibodies per antigen in the SAbDab-filtered CALM dataset for 1,334 unique antigens based on UniProt ID. The inset shows the top ten antigens with the highest number of associated antibodies.

